## Supplementary Information for "Transcription factor recruitment by parallel G-quadruplexes to promote transcription: the case of herpes simplex virus-1 ICP4"

### Table S1. Oligonucleotides used in this study

| Assay | Name | Sequence (5'-3') | 260/290 | $T_m$ (°C) |
|---|---|---|---|---|
| CD | un2 | GGGGGCGAGGGGCGGGAGGGGGCGAGGGG | -0.33 | >90 |
| | un2L2 | GGGGGCGAGGGGCGGGAGGGGGCGAGGGGGGGGG CGAGGGGCGGGAGGGGGCGAGGGG | 11.20 | >90 |
| | myc | GGGGAGGGTGGGGAGGGTGGGGAAGG | 18.66 | >90 |
| | LTR-III | GGGAGGCGTGGCCTGGGCGGGACTGGGG | 0.78 | 65.5 ± 0.1 |
| | hTel | GGGTTAGGGTTAGGGTTAGGG | 0.24 | 68 ± 0.1 |
| | ICP4-146532 | GGGCGGGGCGCGAGGGCGGGTGGG | 3.20 | 79.5 ± 0.1 |
| | ICP4-146666 | GGGGTGGGCCCGCCGGGGGGGCGGGGGG | 8.71 | >90 |
| | ICP4-146574-8 | GGGCGGGGCCGGGGGTTCGACCAACGGGCCGCGGC CACGGG | 2.09 | 76.02 ± 1.53 |
| | ICP4-146947 | GGCGGGGGTCGTCGGGGTCCGTGGG | 5.30 | 69.2 ± 0.2 |
| FRET | F-un2-T | FAM-GGGGGCGAGGGGCGGGAGGGGGCGAGGGG-TAMRA | 25.04 | 74.41 ± 5.10 |
| | F-myc-T | FAM-GGGGAGGGTGGGGAGGGTGGGGAAGG-TAMRA | 52.19 | 78.8 ± 12.45 |
| | F-LTR-III-T | FAM-GGGAGGCGTGGCCTGGGCGGGACTGGGG-TAMRA | 1.25 | 65.80 ± 1.23 |
| | F-hTel-T | FAM-GGGTTAGGGTTAGGGTTAGGG-TAMRA | 0.26 | 67.38 ± 1.05 |
| | un2 complementary | CCCCTCGCCCCCTCCCGCCCCTCGCCCCC | nd | nd |
| | myc complementary | CCTTCCCCACCCTCCCCACCCTCCCC | nd | nd |
| | LTR-III complementary | CCCCAGTCCCGCCCAGGCCACGCCTCCC | nd | nd |
| | hTel complementary | CCCTAACCCTAACCCTAACCC | nd | nd |
| Pull-down/MS XL pull-down/CD | B-un2 | Btn-TTTTTGGGGGCGAGGGGCGGGAGGGGGCGAGGGG | 8.32 | >90 |
| | B-un2L2 | Btn-TTTTTGGGGGCGAGGGGCGGGAGGGGGCGAGGGGG GGGGCGAGGGGCGGGAGGGGGCGAGGGG | 10.92 | >90 |
| | B-myc | Btn-TTTTTGGGGAGGGTGGGGAGGGTGGGGAAGG | 13.67 | >90 |
| | B-gp054dL3 | Btn-TTTTTGGGGTTGGGGCTGGGGTTGGGGGGGGTTGGG GCTGGGGTTGGGGGGGGTTGGGGCTGGGGTTGGGG | 3.12 | >90 |
| | B-LTR-III | Btn-GGGAGGCGTGGCCTGGGCGGGACTGGGG | 0.84 | 68.80 ± 0.77 |
| | B-LTR-II+III+IV | Btn-TTTTTGGGGACTTTCCAGGGAGGCGTGGCCTGGGCGG GACTGGGGAGTGG | 1.39 | 63.07 ± 0.79 |
| | B-ICP4-146532 | BtnTg-GGGCGGGGCGCGAGGGCGGGTGGG | 59.56 | >90 |
| | B-ICP4-146666 | BtnTg-GGGGTGGGCCCGCCGGGGGGGCGGGGGG | 73.11 | >90 |
| | B-ICP4-146574-8 | BtnTg-GGGCGGGGCCGGGGGTTCGACCAACGGGCCGCGGC CACGGG | 11.44 | 72.16 ± 1.182 |
| | B-ICP4-146947 | BtnTg-GGCGGGGGTCGTCGGGGTCCGTGGG | 10.43 | >90 |
| | B-IE3 | Btn-TTTTTCCGATCGTCCACACGGAGC | 0.42 | nd |
| | B-G-rich scrambled | Btn-TTTTTGGAGTCGTGTCGCGTGTCGAGCGTGTGTAGTG GTTTTT | -0.03 | nd |
| FISH; PLA | B-un2 shifted-PLA | Btn Tg-GTTTATTTCGAGGGGCGG | nd | nd |

$T_m$: melting temperatures (°C), calculated from CD thermal unfolding experiments. Nd = not detected. Btn: Biotin, BtnTg: Biotin TEG, FAM: 6-carboxyfluorescein, TAMRA: 6-carboxy-tetramethylrhodamine

**Table S2. Proteins recovered in the pull-down/MS analysis with the four G4 baits**

| | | | | | Score/Match | | | | | | | |
|---|---|---|---|---|---|---|---|---|---|---|---|---|
| | | | | | 8 h.p.i. | | | | 16 h.p.i. | | | |
| | | | | | Elution 1 | | Elution 2 | | Elution 1 | | Elution 2 | |
| **G4 bait** | **Origin** | **Protein acronym** | **Gene name** | **Protein match** | **G4** | **G-rich** | **G4** | **G-rich** | **G4** | **G-rich** | **G4** | **G-rich** |
| **un2L2** | Viral | PAP | UL42 | DNA polymerase processivity factor | 212/16 | ND | 280/17 | 69/13 | 331/10 | ND | 191/9 | 160/6 |
| | Viral | ICP4 | ICP4 | Infected Cell Polypeptide 4 | 186/25 | ND | 235/18 | ND | 196/11 | ND | 97/5 | ND |
| | Viral | MCP | UL19 | Major capsid protein | ND | ND | ND | ND | 277/14 | 67 | 240/13 | ND |
| | Viral | dUTPase | DUT | Deoxyuridine 5'-triphosphate nucleotidohydrolase | 99/13 | ND | 32/5 | ND | ND | ND | ND | ND |
| | Viral | TK | UL23 | Thymidine kinase | 96/7 | ND | ND | ND | 82/2 | ND | ND | ND |
| | Cellular | hnRNP A1L2 | HNRNPA1 L2 | Heterogeneous nuclear ribonucleoprotein A1 like 2 | ND | ND | 41/9 | ND | ND | ND | ND | ND |
| | Viral | TRX-1 | UL38 | Triplex capsid protein-1 | ND | ND | ND | ND | ND | ND | 59/2 | ND |
| | Viral | TRX-2 | UL18 | Triplex capsid protein-2 | ND | ND | ND | ND | ND | ND | 36/4 | ND |
| | Cellular | C23 | NCL | Nucleolin | 31/9 | ND | ND | ND | 151/6 | 114/4 | ND | ND |
| **gp054dL3** | Viral | ICP4 | ICP4 | Infected Cell Polypeptide 4 | 87/4 | ND | 36/2 | ND | | | | |
| | Viral | PAP | UL42 | DNA polymerase processivity factor | 64/4 | ND | ND | ND | | | | |
| | Viral | MCP, VP5 | UL19 | Major capsid protein | ND | ND | 35/2 | ND | | | | |
| **LTR-III** | Viral | ICP4 | ICP4 | Infected Cell Polypeptide 4 | 45/3 | ND | ND | ND | | | | |
| **myc** | Viral | TK | UL23 | Thymidine kinase | 112/3 | ND | ND | ND | | | | |
| | Viral | ICP4 | ICP4 | Infected Cell Polypeptide 4 | 97/5 | ND | ND | ND | | | | |
| | Viral | PAP | UL42 | DNA polymerase processivity factor | 75/6 | ND | 42/3 | 48/3 | | | | |
| | Viral | MCP, VP5 | UL19 | Major capsid protein | ND | ND | 72/4 | ND | | | | |

Protein matches were obtained in two independent experiments. The indicated oligonucleotides were used as G4 baits, while a G-rich oligonucleotide unable to fold into G4 (G-rich scrambled in Table S1) was used as control. Protein hits with scores lower than 30 were not retained, nor those displaying scores higher than 30 in the interaction with the magnetic streptavidin-coated matrix. The two displayed numbers were assigned by Mascot software: Score indicates the probability that the observed match is not a random event; Match indicates the number of fragments that match the recognized protein. ND not detected.

### Table S3. FRET analysis

| Oligonucleotides and proteins | E | ΔE | R (Å) | % unfolding (ΔE) | % unfolding (R) |
|---|---|---|---|---|---|
| **un2** | **0.66±0.003** | **0.30±0.017** | **44.66±0.122** | **0** | **0** |
| ds un2 | 0.36±0.020 | 0 | 55.14±0.801 | 100 | 100 |
| un2 + ipICP4 (5x) | 0.62±0.022 | 0.04±0.002 | 45.94±0.071 | 13.33 | 16.6 |
| un2 + ipICP4 (10x) | 0.58±0.008 | 0.08±0.012 | 47.40±0.261 | 26.66 | 26.14 |
| un2 + ufICP4 (10x) | 0.32±0.003 | 0.34±0.001 | 56.60±0.132 | 114.28 | 102.64 |
| un2 + ipICP4 (5x) + complementary strand | 0.61±0.001 | 0.05±0.018 | 46.18±0.055 | 16.66 | 14.50 |
| un2 + ipICP4 (10x) + complementary strand | 0.52±0.001 | 0.14±0.018 | 49.02±0.015 | 46.66 | 41.60 |
| un2 + ufICP4 (10x) + complementary strand | 0.31±0.002 | 0.35±0.001 | 56.70±0.115 | 113.33 | 102.80 |
| un2 + BSA (10x) | 0.66±0.005 | 0 | 44.61±0.128 | 0 | 0.47 |
| un2 + complementary strand | 0.63±0.090 | 0.03±0.070 | 45.54±0.081 | 10 | 8.39 |
| un2 + BSA + complementary strand | 0.64±0.011 | 0.02±0.008 | 45.29±0.036 | 6.66 | 5.7 |
| un2 + B19 | 1.01±0.007 | -0.35±0.004 | 37.51±0.229 | -53.03 | - 16 |
| un2 + ufICP4 (10x) + B19 | 0.40±0.008 | 0.16±0.005 | 47.41±0.283 | 47.05 | 26.24 |
| **myc** | **0.60±0.008** | **0.22±0.008** | **46.63±0.004** | **0** | **0** |
| ds myc | 0.38±0.009 | 0 | 54.55±0.180 | 100 | 100 |
| myc + ipICP4 (10x) | 0.18±0.003 | 0.42±0.006 | 63.80±0.238 | 190 | 116.95 |
| myc + ufICP4 (10x) | 0.12±0.002 | 0.48±0.007 | 68.96±0.260 | 218 | 126.4 |
| myc + ipICP4 (10x) + complementary strand | 0.13±0.019 | 0.47±0.010 | 68.29±0.198 | 213 | 125.1 |
| myc + pICP4 (10x) + complementary | 0.11±0.001 | 0.49±0.008 | 70.32±0.002 | 222 | 128.9 |
| myc + BSA (10x) | 0.62±0.001 | 0.02±0.007 | 45.89±0.017 | 9 | 9.81 |
| myc + complementary strand | 0.57±0.006 | 0.03±0.003 | 47.51±0.210 | 13.6 | 11.67 |
| myc + BSA + complementary strand | 0.57±0.003 | 0.03±0.006 | 47.61±0.099 | 13.6 | 12.99 |
| **LTR-III** | **0.83±0.005** | **0.82±0.004** | **38.34±0.026** | **0** | **0** |
| ds LTR-III | 0.01±0.001 | 0 | 102.56±0.298 | 100 | 100 |
| LTR-III + ipICP4 (10x) | 0.86±0.004 | 0.03±0.001 | 39.27±0.175 | 3.6 | 2.4 |
| LTR-III + ufICP4 (10x) | 0.82±0.003 | 0.01±0.002 | 38.47±0.178 | 1.21 | 0.2 |
| **hTel** | **0.82±0.003** | **0.81±0.001** | **38.52±0.150** | **0** | **0** |
| hTel + ufICP4 (10x) | 080±0.002 | 0.02±0.002 | 39.36±0.924 | 2.1 | 2.4 |
| ds hTel | 0.004±0.002 | 0 | 126.53±0.937 | 100 | 100 |
| hTel + ipICP4 (10x) | 0.79±0.003 | 0.03±0.001 | 39.73±0.163 | 3.6 | 3.1 |
| hTel + BSA (r=10) | 0.82±0.003 | 0 | 38.60±0.145 | 0.2 | 0 |
| hTel + pICP4 (10x) | 080±0.002 | 0.02±0.001 | 39.36±0.924 | 2.1 | 2.4 |

Energy Transfer (E), Energy Transfer Difference (ΔE) and Radius (Å) of G4-folding oligonucleotides were calculated in the absence/presence of ICP4/BSA proteins and complementary strand. The percentage of unfolding was calculated on the ΔE and R values of the ds oligonucleotide. ipICP4 = immunopurified ICP4; ufICP4 = ultrafiltrated ICP4; BSA = Bovine Serum Albumin; ds = double-stranded.

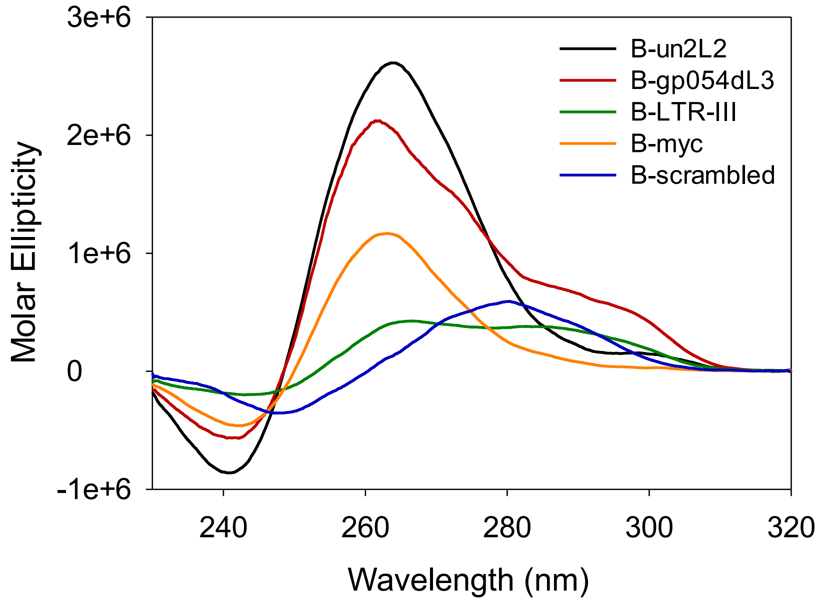


**Figure S1. CD spectra of the biotinylated oligonucleotides used in the pull-down/MS assay.** Oligonucleotides were folded into G4 in potassium phosphate buffer (20 mM PB, 80 mM KCl).

**A**

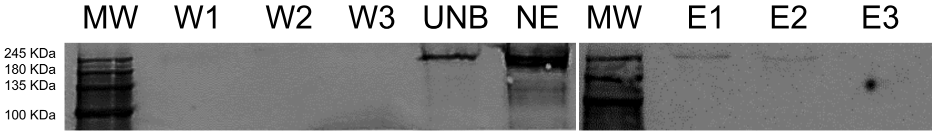


**B**

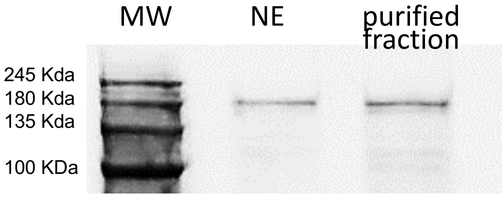


**C**

| Protein | Mass (Da) | Match | Score |
|---|---|---|---|
| Filamin-A OS=Homo sapiens | 283301 | 2/2 | 35 |
| Major viral transcription factor ICP4 OS=Human herpesvirus 1 (strain 17) | 133618 | 13/13 | 136 |
| Transitional endoplasmic reticulum ATPase OS=Homo sapiens | 89950 | 1/1 | 31 |

**Figure S2. ICP4 purification from infected cells. (A)** Western blot of the immunoprecipitated ICP4 (ipICP4) using a monoclonal anti-ICP4 antibody. MW, molecular weight marker IV (Applichem); W1-3, high stringency washes (25 mM Tris-HCl pH 7.4, 150 mM NaCl, 0.5% NP-40, 1 mM EDTA, 5% glycerol); UNB, unbound fraction; NE, nuclear extract (5 µg); E1-3, pH-driven ICP4 elution fractions. **(B)** Western blot of the ultrafiltered ICP4 (ufICP4) from total nuclear extracts (NE) of infected U-2 OS cells (HSV-1 MOI 2). MW is the the molecular weight marker (Applichem, Protein Marker VI (10 – 245 KDa) **(C)** MS analysis of uflICP4 fraction obtained in (B).

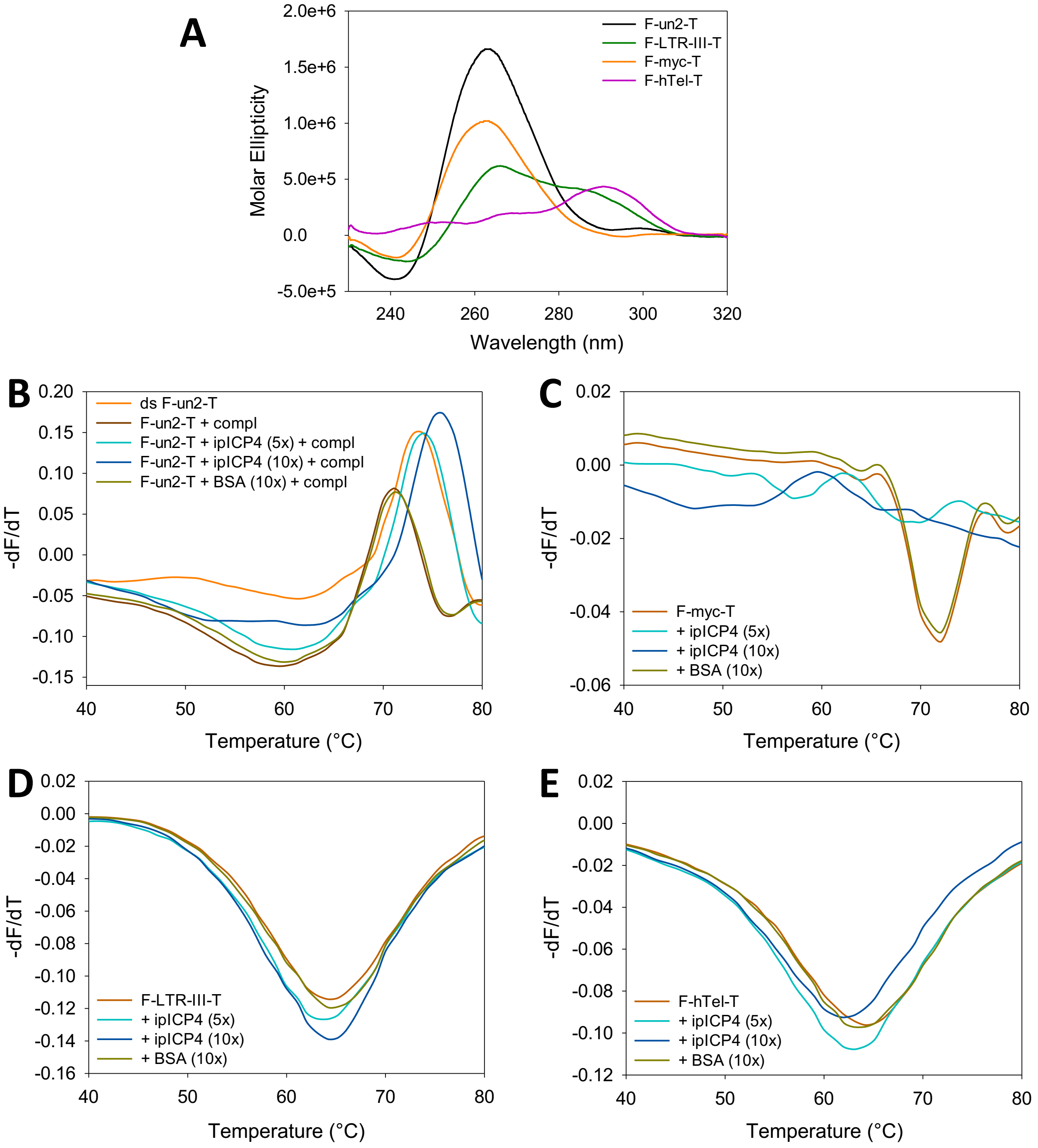


**Figure S3. FRET melting. (A)** CD spectra of the 5'-FAM/3'-TAMRA oligonucleotides used in the FRET melting experiments. The indicated labeled G4 oligonucleotides were folded in potassium phosphate buffer (un2 and myc 20 mM PB, 2 mM KCl; LTR-III and hTel 20mM PB, 80 mM KCl). Different KCl concentrations were used to reach the thermal denaturation of all tested oligonucleotides. **(B-E)** FRET melting analysis of ICP4-mediated unfolding of G4-folded oligonucleotides. **(B)** First derivative FRET-melting curves ($-dF_{525}/dT$ versus $T$) of the folded F-un2-T after incubation with both its complementary strand and ipICP4 (or BSA) at various protein/DNA ratios. The complementary strand and the protein were added at the same time to the G4-folded overnight and incubated for 30 min at 4°C before melting analysis. The complementary strand was used at 1:1 ratio with the G4-folded oligo. As control, un2 denatured and annealed to its complementary strand (1:1 ratio) overnight to yield the full ds oligonucleotide was used. **(C-E)** First derivative FRET-melting curves ($-dF_{525}/dT$ versus $T$) of the indicated G4 oligonucleotides treated with ipICP4 or BSA as control at various protein/DNA ratios, in 20 mM PB pH 7.4 and 2-80 mM KCl.

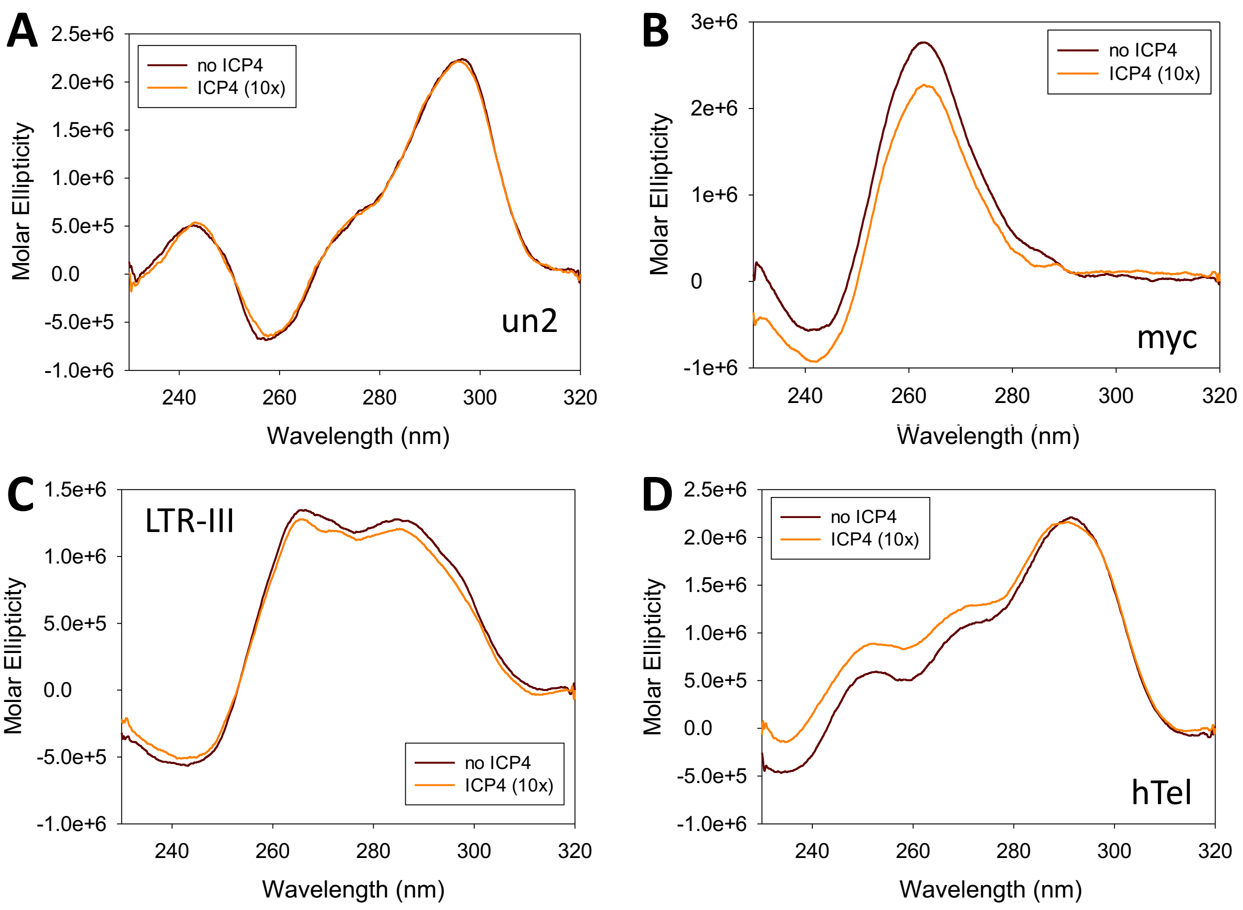


**Figure S4. CD spectra in the presence/absence of ufICP4.** The indicated unlabeled G4 oligonucleotides were folded in potassium phosphate buffer (un2 and myc 20 mM PB, 2 mM KCl; LTR-III and hTel 20mM PB, 80 mM KCl) and incubated in the absence/presence of ufICP4 (10x protein/DNA ratio). Spectra of ICP4 alone was subtracted from G4-ICP4 complex spectra.

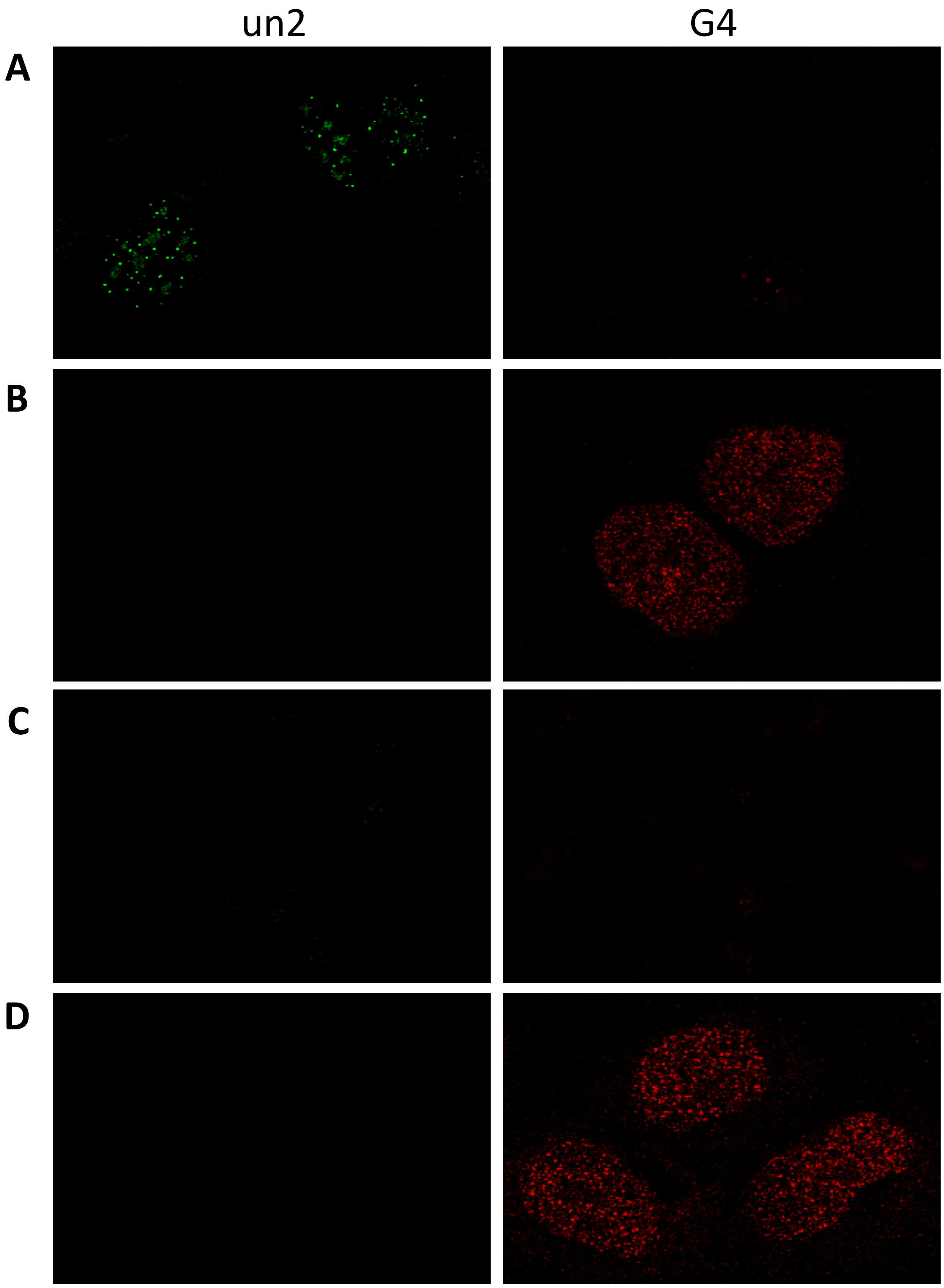


**Figure S5. Controls of immuno FISH analysis.** U-2 OS cells were infected with HSV-1, strain F (MOI 2,) and fixed at 8 hpi. **(A)** Control lacking 1H6 Ab for the detection of G4s; un2 probe and secondary Abs for the un2 probe and 1H6 Ab are present. **(B)** Control lacking the un2 probe; 1H6 Ab and secondary Abs for the un2 probe and 1H6 Ab are present. **(C)** Mock sample (non-infected U-2 OS cell) lacking 1H6 Ab; un2 probe and secondary Abs for the un2 probe and 1H6 Ab are present. **(D)** Mock sample (non-infected U-2 OS cell) lacking un2 probe; 1H6 Ab and secondary Abs for the un2 probe and 1H6 Ab are present.

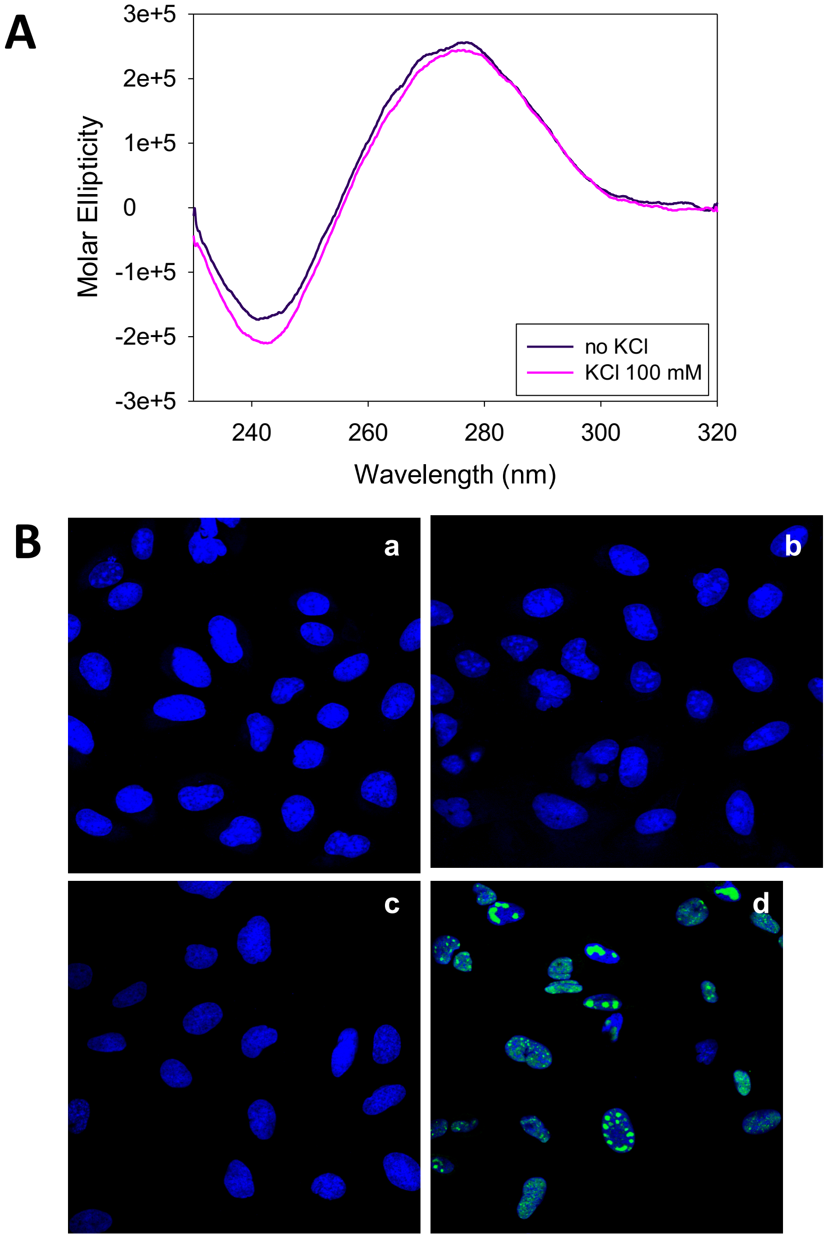


**Figure S6. Controls of PLA analysis. (A)** CD spectra of the un2 PLA probe that does not display a G4 signature. **(B)** PLA controls on mock (uninfected) U-2 OS cells (merged signals, blue staining detects nuclear nucleic acids): (a) cells incubated with un2 probe and subjected to the complete PLA procedure; (b) cells incubated in the absence of un2 probe and subjected to the complete PLA procedure; (c) cells incubated with un2 probe in the absence of primary antibodies and subjected to the complete PLA procedure. No false positive/unspecific PLA dots were observed; (d) U-2 OS infected cells (MOI = 3) processed for probe annealing (high temperature step) and incubated with the mouse anti-ICP4 primary antibody followed by the FITC-labeled secondary antibody (green signal) as control of the primary antibody ICP4 recognition in cells processed for PLA assay.

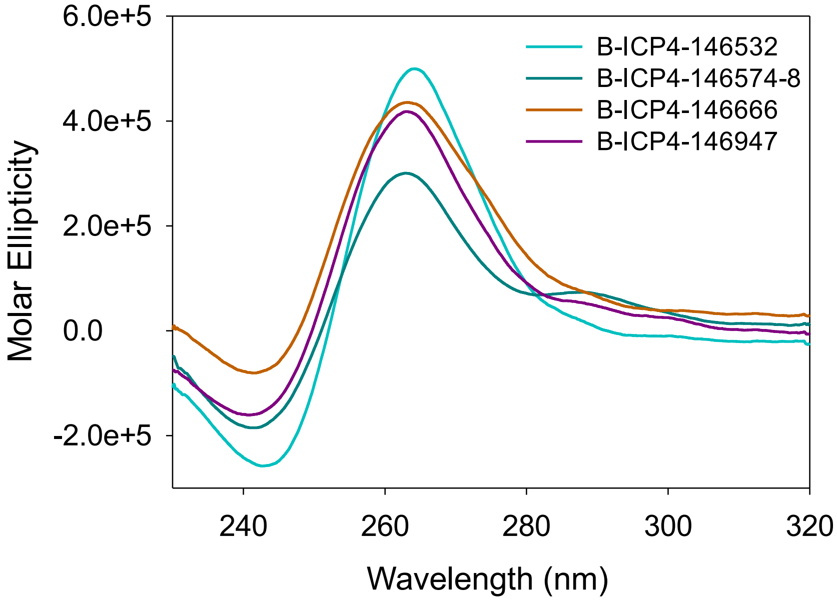


**Figure S7. CD spectra of the biotinylated G4-forming oligonucleotides present in the ICP4 promoter.** The oligonucleotides were folded in potassium phosphate buffer (20 mM PB, 80 mM KCl).
